# Automated Electrodes Detection during simultaneous EEG/fMRI

**DOI:** 10.1101/395806

**Authors:** Mathis Fleury, Christian Barillot, Marsel Mano, Elise Bannier, Pierre Maurel

## Abstract

The coupling of Electroencephalography (EEG) and functional magnetic resonance imaging (fMRI) enables the measure of brain activity at high spatial and temporal resolution. The localisation of EEG sources depends on several parameters including the knowledge of the position of the electrodes on the scalp. An accurate knowledge about this information is important for source reconstruction. Currently, when acquiring EEG and fMRI together, the position of the electrodes is generally estimated according to fiducial points by using a template. In the context of simultaneous EEG/fMRI acquisition, a natural idea is to use magnetic resonance (MR) images to localise EEG electrodes. However, most MR compatible electrodes are built to be almost invisible on MR Images. Taking advantage of a recently proposed Ultra short Echo Time (UTE) sequence, we introduce a fully automatic method to detect and label those electrodes in MR images. Our method was tested on 8 subjects wearing a 64-channel EEG cap. This automated method showed an average detection accuracy of 94% and the average position error was 3.1 mm. These results suggest that the proposed method has potential for determining the position of the electrodes during simultaneous EEG/fMRI acquisition with a very light cost procedure.

## 2 Introduction

Electroencephalography (EEG) measures the electrical potential generated by the neuronal activity over the scalp with electrodes placed on the surface of the scalp (Buzsáki et al. [2012], Murakami and Okada [2006], Petsche et al. [1984]). Usually electrodes are placed thanks to a flexible cap and positioned according to anatomical points enabling optimal covering of brain regions regardless of the size and shape of the subject’s head. Currently, when acquiring EEG and functional magnetic resonance imaging (fMRI) simultaneously, the position of the electrodes is calculated according to fiducial points (anatomical points of the skull) such as inion, nasion and vertex (Strobel et al. [2008]). The localisation of EEG sources in the brain depends on several parameters including the position of the electrodes on the scalp. A precise knowledge of these positions is important because inaccurate information on EEG electrodes coordinates may affect EEG inverse solution (Khosla et al. [1999]). This knowledge is even more crucial in the case of simultaneous EEG and fMRI study, when the sessions are conducted repeatedly over a long period of time. Approximations in the positioning of the electrodes are then made in each session and will give rise to important inaccuracies in the measured evoked potential (Wood and Allison [1981]). As a matter of fact, magnetic resonance (MR) images and EEG need to be registered to be able to compare activations given by fMRI and by EEG. This simultaneous acquisition allows the concordance of two different kind of information, a high temporal resolution in the order of a millisecond with EEG, and a high spatial resolution in the order of millimetre with MRI.

In this article an automated and efficient method to determine EEG electrodes positions based on a specific MR sequence is presented and evaluated. Compared to other existing approaches, the proposed method does not need additional hardware (like 3D electromagnetic digitizer devices (Adjamian et al. [2004];Whalen et al. [2008]), artificial electrode markers (Sijbers et al. [2000]) or laser scanner(Koessler et al. [2011]; Bardouille et al. [2012])), which might be uncomfortable for the subject if he must stay still during acquisition (Le et al. [1998]) and add time to the preparation of the patient. Semi-automated electrodes localisation methods exist (de Munck et al. [2012]; Butler et al. [2017]), which require a manual fiducial landmark identification to guide co-registration without any markers but these approach relies on the efficiency of the accuracy of the operator. Another automated method was recently developed and shown great results with an anatomical MR image (Marino et al. [2016]), however, this method is only working with a high density cap also compatible with MRI: the GES 300 from Geodesic EEG Systems. Since this kind of cap includes plastic around electrodes and contain hydrogen protons, it can be visible on T1-w image. For seek of genericity (i.e. able to operate on all types of caps when artefacts do not appear on T1-w images), we propose to make use of a MRI sequence with radial k-space sampling named UTE for Ultra-short Echo-Time. It allows to visualise the tissues with a very short T2 and T2*, such as cortical bone, tendons and ligaments (Holmes and Bydder [2005], Keereman et al. [2010]). This sequence is all the more interesting in our context because it enables the visualisation of the MR compatible electrodes (Butler et al. [2017], Springer et al. [2008]) on the scalp with a capability to be performed rapidly enough to not overwhelm the whole MRI protocol.

We propose a fully automated method, which provides reliable and reproducible results for the detection and labelling of a MR compatible EEG cap into the MR space.

## 3 Methods

The retrieval of the electrodes consisted in two parts; firstly, we provided a mask that includes the volume where the electrodes are located; secondly, we performed the electrode detection inside this volume of interest (VOI). Figure 1 presents a flowchart of the method’s main steps. We hypothesised that electrodes would appear as spheres inside the UTE volume and it allows us to perform a Hough transform in a consistent manner across subjects.

**Figure 1:**
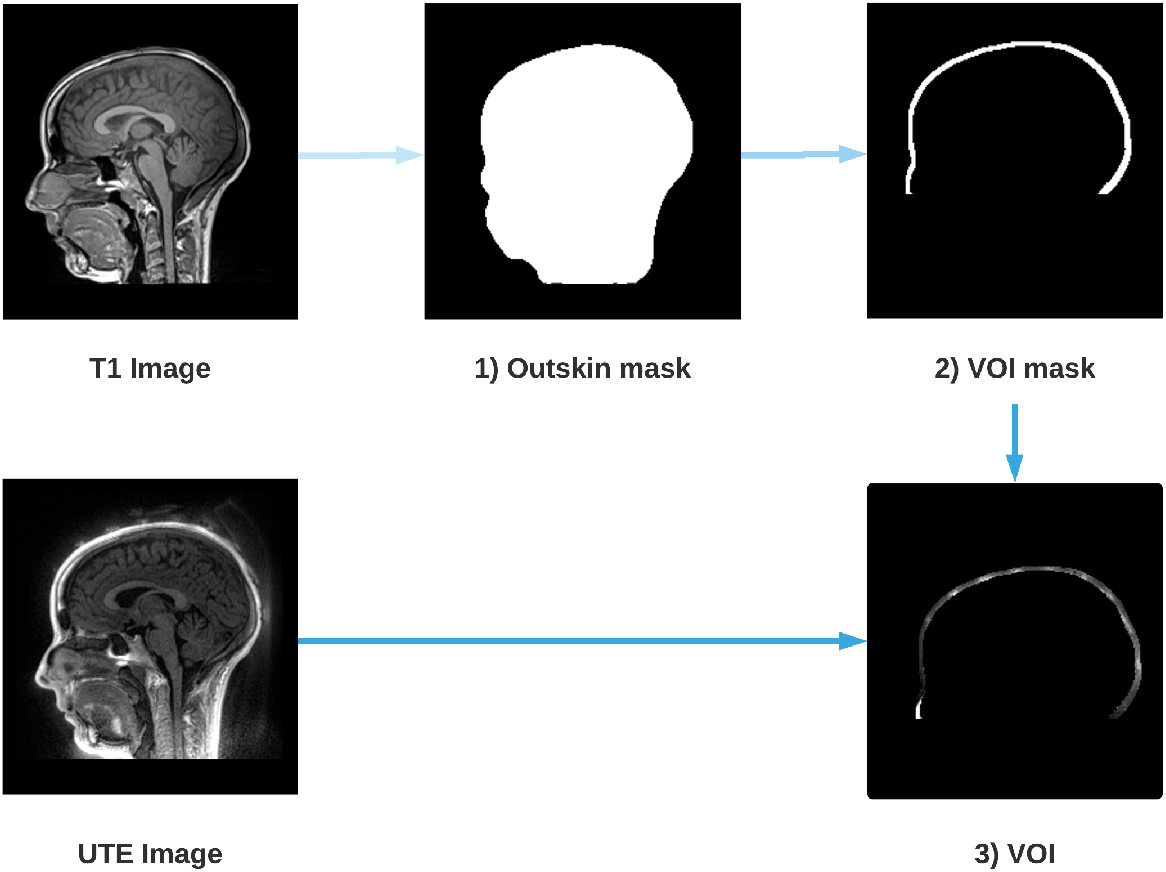
Steps for the extraction of the Volume Of Interest (VOI). An outskin mask is performed from the T1 image (1), then a dilation and a removal of the mask is performed (2) in order to obtain the layer where the electrodes are located. Finally, the UTE image is masked by the dilated mask (2) which gives us the VOI (3).

### 3.1 Scalp segmentation

Several reliable scalp segmentation methods exist for T1-w imaging. Because UTE images are noisier, we performed the scalp segmentation on the T1-w images and co-registered the UTE images with the T1-w images to apply the mask. The T1-w is first registered on the UTE and the anatomical T1 image is then segmented using FSL, an open-library of analysis tools for MRI and its function BET (Brain Extraction Tool) (Smith [2002]; Popescu et al. [2012]). A mask of the scalp is computed from the segmentation. Since electrodes are located around the head of the subject, the scalp mask is dilated toward the periphery in order to isolate this layer. What is outside the dilated mask is subtracted in order to isolate only the layer where the electrodes are located.

### 3.2 Detection of electrodes with the Spherical Hough transform

A 3D Hough transform was used to segment the electrodes inside the VOI. Hough transform is typically used to detect circles or lines in 2-dimensional data sets, but was recently extended to detect spheres in 3-dimensional data sets (Xie et al. [2012]; Borrmann et al. [2011]). As the shape of an electrode can be assimilated to a sphere, the Spherical Hough Transformation algorithm seemed particularly well adapted to this task. The VOI image is first smoothed using a Gaussian kernel, with a FWHM (Full Width at Half Maximum) adapted to the size of the electrode (10 mm) in order to reduce the noise of the image while saving electrode information. Then, the Hough algorithm is performed and provides a list of n potential electrodes, *D* = [*d*_1_,…, *d_n_*]. Figure 2 shows an example of such detections on a 2D slice of the VOI. Because the VOI includes also anatomical structures (nose, ears) and noise (artefacts due to the cap or gel), the number of potentially detected electrodes is substantially higher than the number of “true” electrodes *N*, in our case 64.

**Figure 2:**
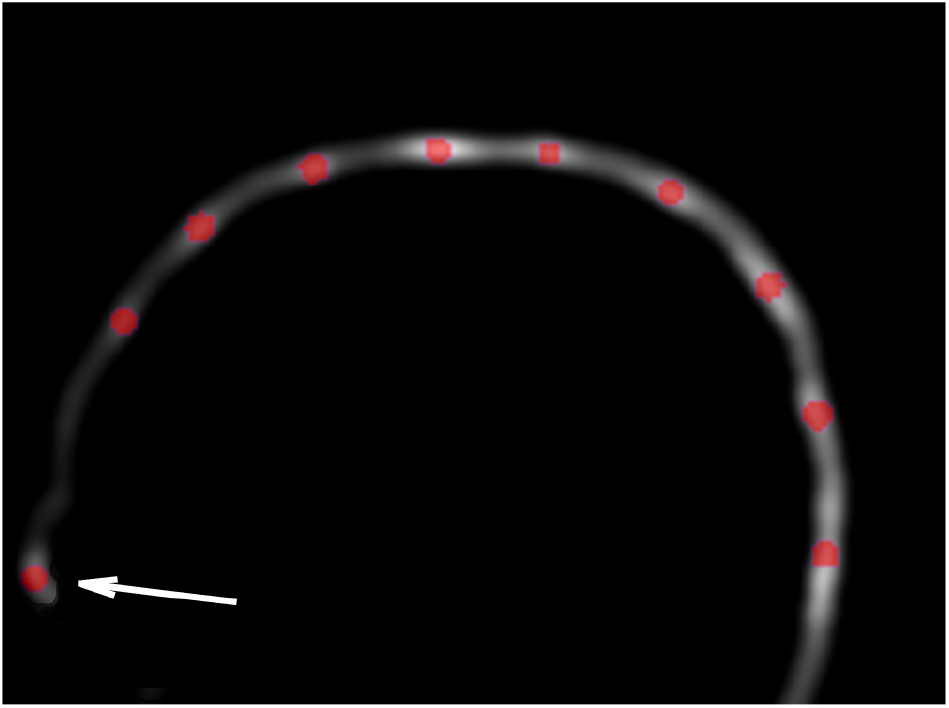
Example of Hough transform detection (red dots) on the VOI smoothed image. Hough transform detects also anatomical parts (arrow), which will be excluded in the filtering steps (cf. section 3.3).

### 3.3 Selection of detected electrodes

The detected electrodes are then filtered to get rid of the potential false detections given by the Hough transform. A 64 electrodes spherical EEG template *p_j_* (1 ≤ *j* ≤ 64) ∈ *P* was given by the cap manufacturer, indicating theoretical positions of every electrodes relatively to each other. Due to the non-sphericity of the head and the elastic deformations of the cap, these positions are not sufficient enough to give a reliable detection by itself. However, this template will be used to identify outliers in our detections. This spherical template is registered onto the detected electrodes from previous section, through the Iterative Closest Point (ICP) algorithm, a well-known algorithm for registering two-cloud of points (Besl et al. [1992]; Chen and Medioni [1992]). The algorithm takes a first point cloud which will be kept fixed, while the other one will be spatially transformed in order to best align the reference. The goal is to iteratively minimise a metric error, usually the distance between the two sets of points, by modifying the transformation applied to the source.

In our case, the ICP will find the optimal rotation, translation and scale to fit the data point set *D* obtained with the Hough transform and the model point *P*. The algorithm is divided into 2 steps. The first step consists in estimating correspondences between the two set of points. During this step, for each point *p_j_*, in the reference set *P*, the closest point *d_i_* of the detected points set *D* is computed. This point will be noted *c_j_* and therefore defined as follows:

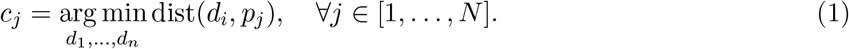

The second step consists in computing the similarity transform that best aligns every *c_j_* to the corresponding *p_j_*. The minimisation is expressed by:

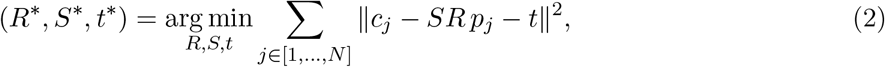

where *R* is a rotation matrix (3 × 3), *t* is a translation vector (3 × 1) and *S* is a scale matrix (*S* = *s * Id*, 3 × 3). The ICP runs until convergence. The registered template *P*′ can then be written as:

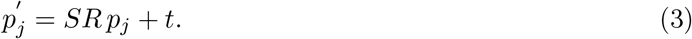

Once the ICP is completed, a two-part filtering phase is implemented. The first one consists in taking the closest point of the Hough transform data set; for each of the *N* electrodes of the registered model *P*′ the closest detected point *c_j_* is selected. Unselected points are discarded and, after this first filtering step, the number of electrodes is therefore equal to *N*, the total number of electrodes desired (64 in our case). Figure 3 illustrates the impact of this step.

**Figure 3:**
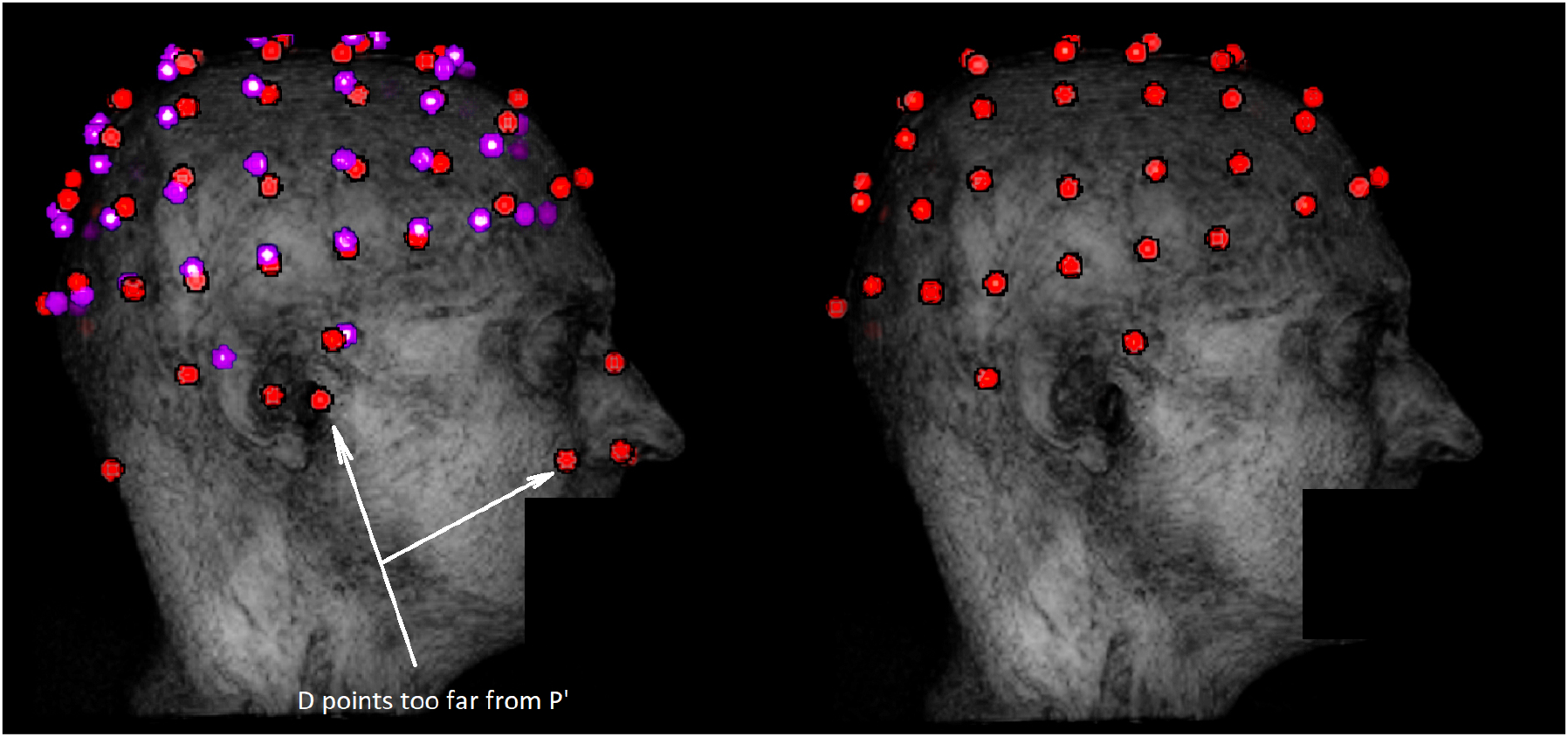
Example of outliers removal in potential electrodes data set *D* with the ICP algorithm. The dataset *D* is represented in red on the left along with the registered template *P*′ in purple. The data set obtained after the first filtering step is in red on the right. Outliers are mostly due to external anatomical parts or noise not taken in account during the segmentation. These outliers are discarded by the filtering step because they are too far from *P*′.

For the second and final step, all points *c_j_*, which are too far from the closest point of the template *P*′, are removed. A threshold equals to four times the Median Absolute Deviation (MAD) of all distances is applied. For each removed point, a replacement is determined by a new detection from the local maxima on the VOI image around the theoretical position given by the registered template (cf. Figure 4). The new data set *D*′ is obtained and the *N* electrodes are then labeled using the template.

**Figure 4:**
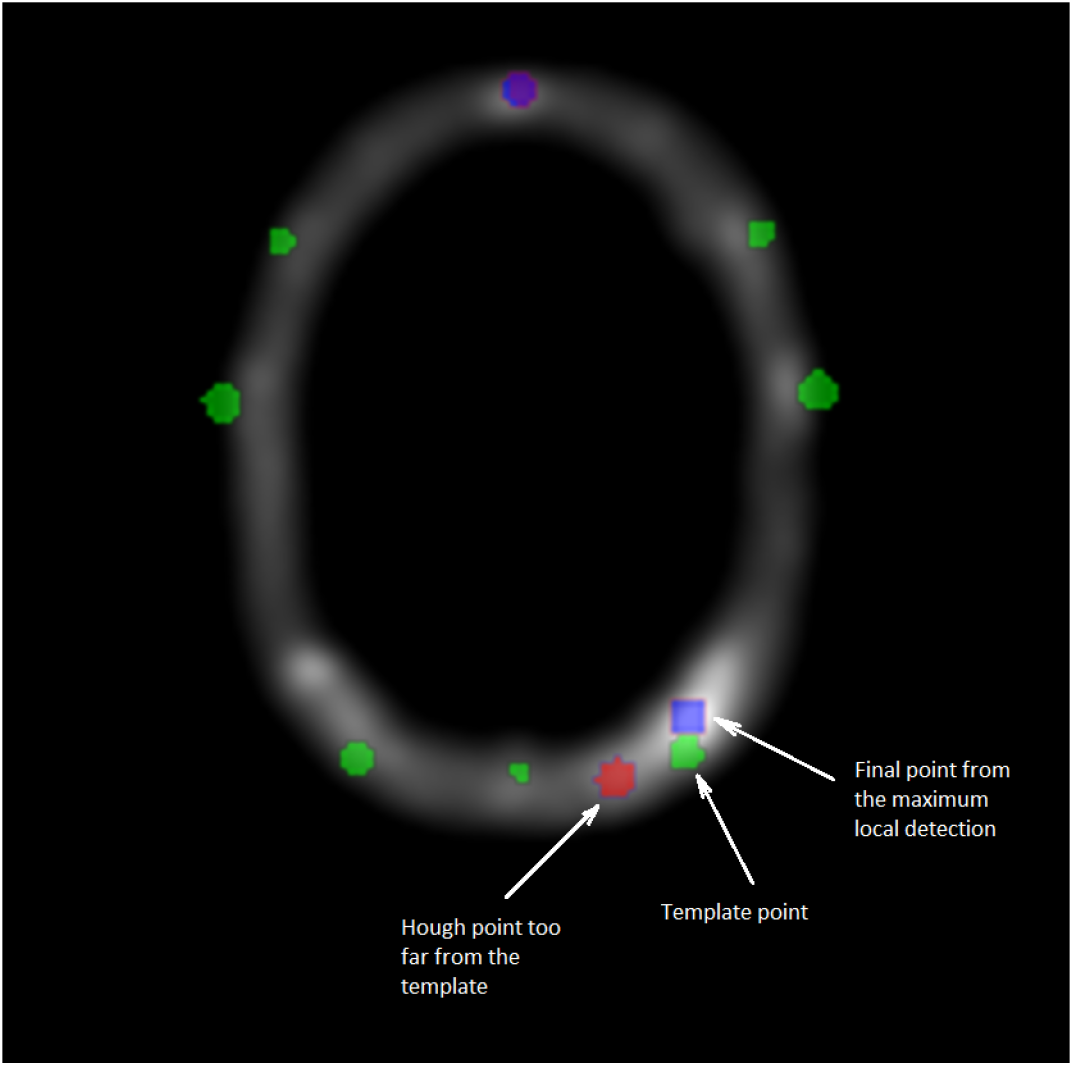
Cross section of the VOI image. Green points are corresponding to the template data set *P*′, blue points to the maximum local detection and the red one are the outliers from *D*. The second and final filtering step consists in replacing any point from the Hough data set too far from the registered template *P*′. The substituted point comes from a detection by local maxima, closest to the template *P*′.

### 3.4 Validation of the method

A manual selection of the electrodes positions was done on the UTE sequence and the quality of our detection was assessed using this manual selection as a ground truth. Instead of selecting the center of each electrode in a 3D image, we choose to use a more convenient procedure for the manual detection. Following Butler et al. [2017], the manual detection was performed by picking up the Cartesian position (*x_i_,y_i_,z_i_*) of each 64 electrodes for each subject on a pancake view, which is roughly a 2D projection of the scalp (de Munck et al. [2012]).

The performance indicators of our automated detection will be the position error (PE) and the positive predictive value (PPV). The position error is the average Euclidean distance between each pair of electrodes (the manually selected one, considered as the ground truth, and the detected one) and the PPV is the percentage of electrodes that have been well detected. We considered that a detected electrodes is well localised when the PE is below 10 mm, which corresponds to the diameter of the electrode (Kavanagk et al. [1978]).

We also compared the performance of our method against a more traditional semi-automatic one: five fiducial points were selected manually and the spherical template was adjusted to these points (Towle et al. [1993]). This method, although not recent, is still used by many studies (e.g. Thornton et al. [2017], Jenson et al. [2018], Ge et al. [2017]).

## 4 Materials

### 4.1 Subjects and EEG equipment

After IRB approval, eight healthy volunteers provided written informed consent to take part in the study. They all underwent a simultaneous EEG/fMRI examination (fully described in Mano et al. [2017]). EEG was acquired using two 32-channel MR compatible amplifiers (actiCHamp, Brainproduct, Gilching, Germany) and a cap providing 64 Ag/AgCl electrodes positioned according to the extended 10-20 system and one additional ground electrode. Electrodes are attached to small cups with inner diameter of 10 mm and 4 mm height, inserted in the cap and filled with gel to minimise the contact impedance. All subject wore a large (circumference between 56-58 cm) MR compatible cap from Brainproduct (Gilching, Germany) and a particular attention was given to its positioning according to standard fiducial points.

### 4.2 UTE sequences parameters

All MR data were collected on a 3T Siemens Verio MR scanner (VB17, Siemens Healthineers, Erlangen, Germany). Specifically, the UTE sequence using 3D radial k-space sampling was performed with the following parameters: repetition time (TR) = 3.45 ms, echo time (TE) = 0.07 ms, flip angle (FA) = 14° and voxel size 1.33 × 1.33 × 1.33 mm^3^. A 3D T1 MPRAGE was also performed: TR = 1900 ms, TI = 900 ms, TE = 2.26 ms, FA = 9° and voxel size 1 × 1 × 1 mm^3^. Two additional UTE sequences with lower sampling resolution were acquired in order to decrease the acquisition time and to investigate the impact on electrodes detection. To reduce the acquisition time, the number of spokes has to decrease; from 60000 spokes (60K) for the original, to 30000 (30K) and 15000 (15K) spokes for the additional ones. The UTE acquisition time goes down from 5 min 35 s to 2 min 47 s and 1 min 23 s. A comparison between these acquisitions is shown in Figure 5.

**Figure 5:**
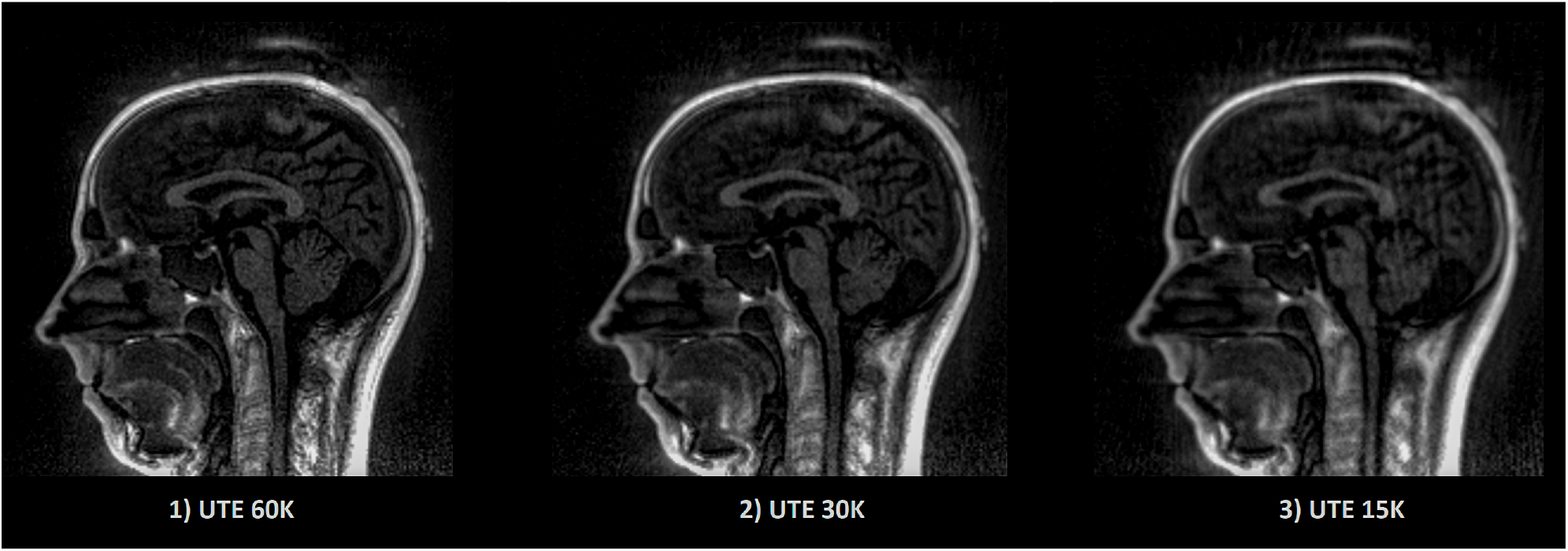
Example of UTE images with different sampling. The image quality as well as the acquisition time decrease linearly according to the sampling. Acquisition time for 1) 5 min 35 s, 2) 2 min 47 s, 3) 1 min 23 s.

## 5 Results

The creation of an image (VOI) containing only the information related to the electrode allows to remove external noise while protecting the information related to the electrode. This image enables robust detection of the position of the electrodes for all subjects. Furthermore, since our method always detects exactly *N* (64 in our case) electrodes, the number of false negatives (missed electrodes) will automatically be equal to the number of false positives (wrongly detected electrodes). Table 1 presents the mean position error (PE), the standard deviation of the PE and the maximum PE of our detections for each of the eight subjects. The max PE reflected a high difficulty to detect the electrodes near anatomical parts or in posterior regions where the head apply a pressure on the EEG cap inside the MRI. Our UTE-based electrode detection showed an average PE of 3.1mm for all subjects. The detection accuracy, represented by the positive predictive value (PPV), is also shown and corresponds to the percentage of electrodes correctly found. The average PPV for all subjects was 94.22%.

**Table 1:**
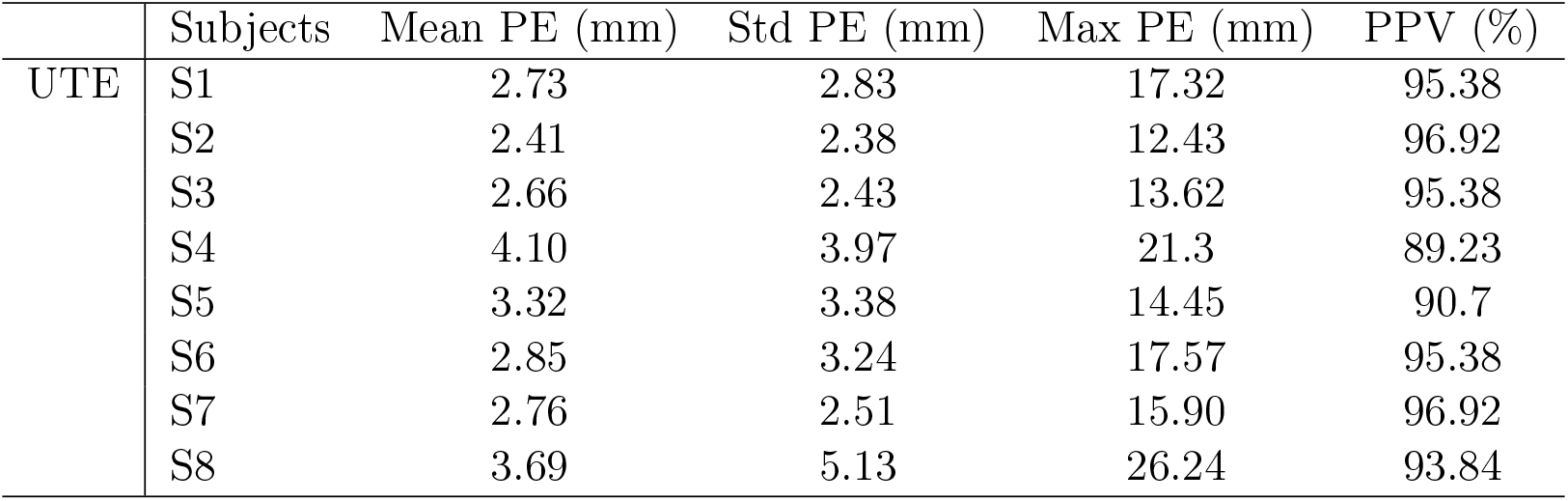
Position error (PE) and positive predictive value (PPV) for each subject (S1-S8) for UTE-MR electrodes detection. The PPV is the percentage of electrodes that have been detected. We consider that an electrode is well localised when the PE is below 10 mm, which represents the diameter of an electrode. The mean PE on all subject is equal to 3.1mm and the mean PPV to 94.22%.

|  | Subjects | Mean PE (mm) | Std PE (mm) | Max PE (mm) | PPV (%) |
| --- | --- | --- | --- | --- | --- |
| UTE | S1 | 2.73 | 2.83 | 17.32 | 95.38 |
|  | S2 | 2.41 | 2.38 | 12.43 | 96.92 |
|  | S3 | 2.66 | 2.43 | 13.62 | 95.38 |
|  | S4 | 4.10 | 3.97 | 21.3 | 89.23 |
|  | S5 | 3.32 | 3.38 | 14.45 | 90.7 |
|  | S6 | 2.85 | 3.24 | 17.57 | 95.38 |
|  | S7 | 2.76 | 2.51 | 15.90 | 96.92 |
|  | S8 | 3.69 | 5.13 | 26.24 | 93.84 |

We then compared the performance of our method with the semi-automatic one presented in section 3.4 (FID). The PE and PPV were calculated in the same way. The results are shown in Table 2 and Figure 6 shows, for each subject, a comparison of the PEs obtained by the two methods. The mean PE on all subject is equal to 7.7 mm and the mean PPV to 79.41%. Moreover, for every subjects, our method produced smaller PE and better PPV. A paired t-test was computed between the two PEs sets and a significant difference was obtained (p<0.0001).

**Figure 6:**
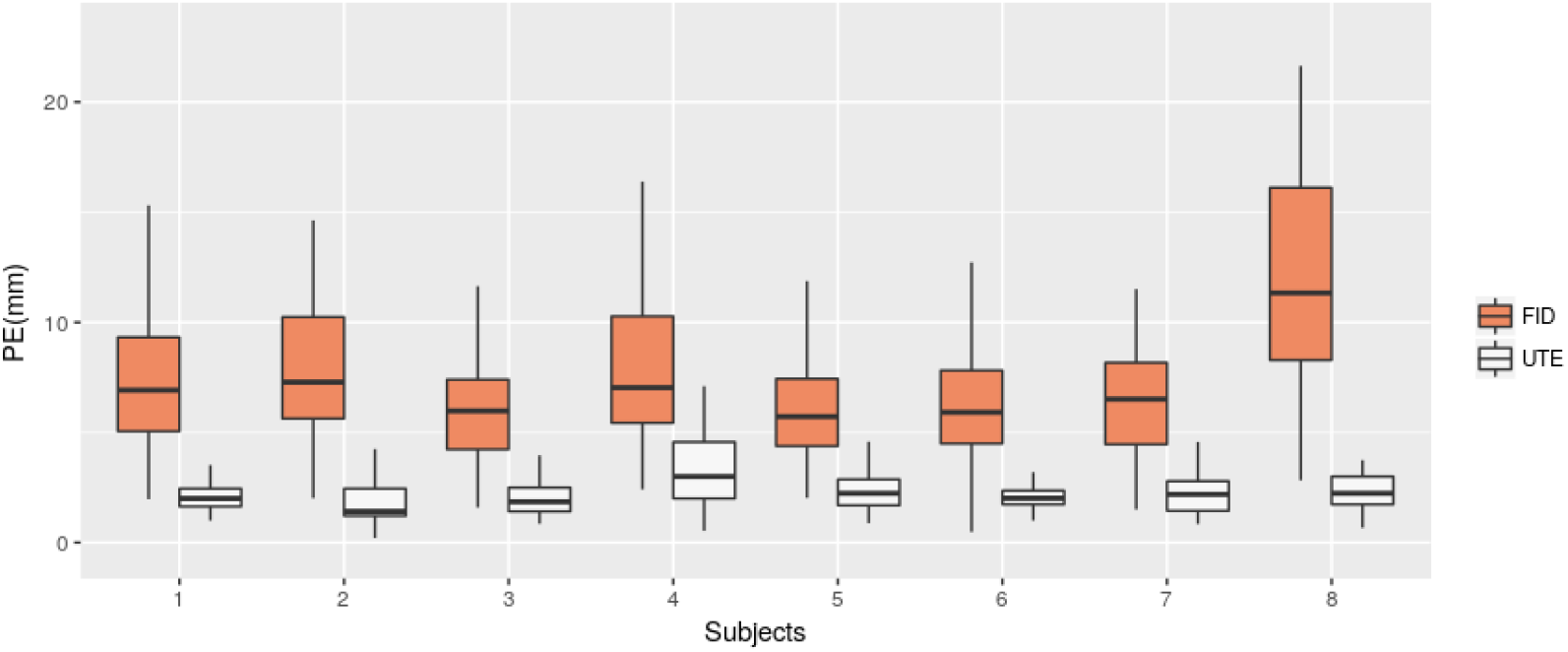
Position Error (PE) for UTE-based electrodes detection method (UTE) and the semi-automatic method based on fiducial points (FID). Box-plots for the eight subjects are shown.

**Table 2:**
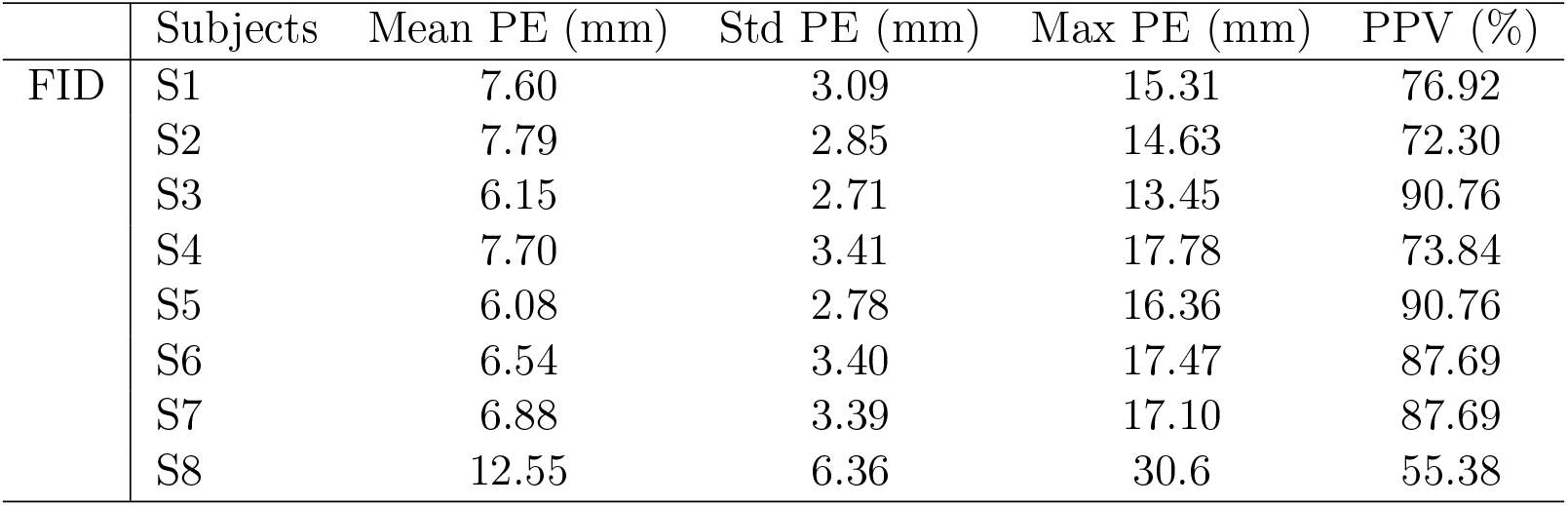
Positive predictive value (PPV) and position error (PE) for each subject (S1-S8) for semi-automated electrodes detection based on manual delineation of fiducial landmark (FID). The PPV is the percentage of electrodes that have been detected. We consider that an electrode is well localised when the PE is below 10 mm, which represents the diameter of an electrode. The mean PE on all subject is equal to 7.7 mm and the mean PPV to 79.41%.

|  | Subjects | Mean PE (mm) | Std PE (mm) | Max PE (mm) | PPV (%) |
| --- | --- | --- | --- | --- | --- |
| FID | S1 | 7.60 | 3.09 | 15.31 | 76.92 |
|  | S2 | 7.79 | 2.85 | 14.63 | 72.30 |
|  | S3 | 6.15 | 2.71 | 13.45 | 90.76 |
|  | S4 | 7.70 | 3.41 | 17.78 | 73.84 |
|  | S5 | 6.08 | 2.78 | 16.36 | 90.76 |
|  | S6 | 6.54 | 3.40 | 17.47 | 87.69 |
|  | S7 | 6.88 | 3.39 | 17.10 | 87.69 |
|  | S8 | 12.55 | 6.36 | 30.6 | 55.38 |

Finally, we investigated the impact of lower sample UTE sequences, which allow reducing the acquisition time, on electrode detection. We tested two others UTE sequence (cf. Section 4.2). We applied our detection method on the three different UTE images and compared the quality of the detections. Table 3 reports the mean PE and mean PPV obtained for the three UTE sequences on seven subjects (the first subject did not receive the additional sequences). As expected, the mislocalisation, as well as the position error, increase according to the decrease of the sampling. However, our results are still clearly better than the semi-automatic one for the 30k sequence (half the acquisition time than the original one) and are slightly better for the fastest sequence.

**Table 3:**
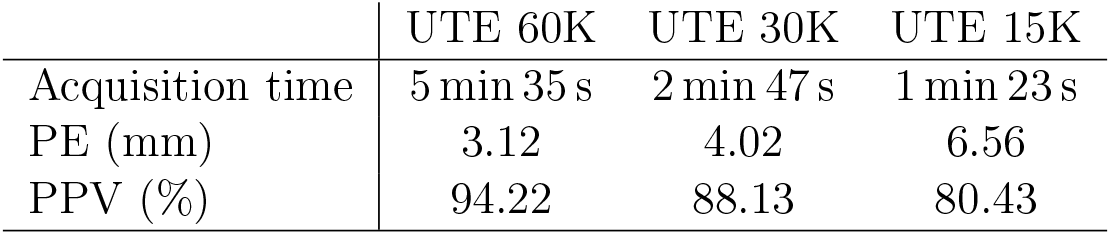
Mean of position error (PE) and mean positive predictive value (PPV) for three different sampling resolutions of the UTE sequence. Shorter acquisition time implied lower SNR and lower detection accuracy. Results are still better than the semi-automatic method.

|  | UTE 60K | UTE 30K | UTE 15K |
| --- | --- | --- | --- |
| Acquisition time | 5 min 35 s | 2 min 47 s | 1 min 23 s |
| PE (mm) | 3.12 | 4.02 | 6.56 |
| PPV (%) | 94.22 | 88.13 | 80.43 |

## 6 Discussion

We have proposed an automated method for detecting and labelling EEG electrodes based on UTE MR images without using any external sensors. Previous results indicate that a localisation technique using electromagnetic digitisation technology is time-consuming (Dalal et al. [2014]) and others techniques such as 3D digitisation can be affected by errors of registration and projection of EEG electrodes on the head model. We have shown that our method offers constant and precise results. Moreover, the proposed method provides the position of the electrodes directly into the MR-space, which is crucial in case of simultaneous EEG/fMRI acquisitions.

Furthermore, for seek of genericity, the proposed method is able to operate on all types of caps and does not need specific electrodes, unlike a recent work from Marino et al. [2016] for example. To the best of our knowledge, this is first automated electrodes detection method implying non-visible electrodes on anatomical MR sequence.

The method presented here requires only an additional sequence (the UTE acquisition sequence) in the experimental protocol. This acquisition takes from 1 to 5 min. From our experiments, a good compromise between acquisition time and detection quality can be achieved with a 2 or 3 min sequences. Further optimisation of the sequence parameters could enable an improvement of the images without increasing the acquisition time.

## 7 Conclusion

We presented a method to automatically detect and label EEG electrodes during an EEG/fMRI acquisition. We used a UTE MR sequence to obtain electrodes positions on a MR-volume. This method only has for additional cost the acquisition time of the UTE sequence in the MR protocol. We have demonstrated that our method achieves a significantly more accurate electrode detection compared to a semi-automatic detection one that is more commonly used during EEG/fMRI protocols. For future research, since the proposed method can be totally automated and does not require complex processing, this technique may be used to extract the position and the label of the electrodes in real time. Indeed, this technique is interesting for applications requiring immediate knowledge of the position of the electrodes. We believe this method will be useful to improve the fusion of EEG and fMRI signals.

## 8 Acknowledgement

MRI data acquisition was partly supported by the Neurinfo MRI research facility from the University of Rennes I. Neurinfo is granted by the the European Union (FEDER), the French State, the Brittany Council, Rennes Metropole, Inria, Inserm and the University Hospital of Rennes. The authors would like to thank their collaborators at Siemens Healthineers, in particular Vladimir Jellus and P. Speier for their valuable contributions to the development and testing of the prototype Ultra Short TE for this project.

